# *Maturity2*, a novel regulator of flowering time in *Sorghum bicolor*, increases expression of *SbPRR37* and *SbCO* in long days delaying flowering

**DOI:** 10.1101/535484

**Authors:** Anna L. Casto, Ashley J. Mattison, Sara N. Olson, Manish Thakran, William L. Rooney, John E. Mullet

**Affiliations:** Biochemistry and Biophysics Department, Texas A&M University, College Station, TX, United States of America; Soil and Crop Science Department, Texas A&M University, College Station, TX, United States of America

## Abstract

*Sorghum bicolor* is a drought-resilient facultative short-day C4 grass that is grown for grain, forage, and biomass. Adaptation of sorghum for grain production in temperate regions resulted in the selection of mutations in *Maturity* loci (*Ma_1_ – Ma_6_*) that reduced photoperiod sensitivity and resulted in earlier flowering in long days. Prior studies identified the genes associated with *Ma_1_* (*PRR37*), *Ma_3_* (*PHYB*), *Ma_5_ (PHYC*) and *Ma_6_ (GHD7*) and characterized their role in the flowering time regulatory pathway. The current study focused on understanding the function and identity of *Ma_2_. Ma_2_* delayed flowering in long days by selectively enhancing the expression of *SbPRR37 (Ma_1_*) and *SbCO*, genes that co-repress the expression of *SbCN12*, a source of florigen. Genetic analysis identified epistatic interactions between *Ma_2_* and *Ma_4_* and located QTL corresponding to *Ma_2_* on SBI02 and *Ma_4_* on SBI10. Positional cloning and whole genome sequencing identified a candidate gene for *Ma_2_*, Sobic.002G302700, which encodes a SET and MYND (SYMD) domain lysine methyltransferase. Nine sorghum genotypes previously identified as recessive for *Ma_2_* contained the mutated version of Sobic.002G302700 present in 80M (*ma_2_*).

## Introduction

*Sorghum bicolor* is a drought resilient, short-day C4 grass that is grown globally for grain, forage and biomass [1–4]. Precise control of flowering time is critical to achieve optimal yields of sorghum crops in specific target production locations/environments. Sorghum genotypes that have delayed flowering in long days due to high photoperiod sensitivity are high-yielding sources of biomass for production of biofuels and specialty bio-products [3,5]. In contrast, grain sorghum was adapted for production in temperate regions by selecting genotypes that have reduced photoperiod sensitivity resulting in earlier flowering and reduced risk of exposure to drought, heat, or cold temperatures during the reproductive phase. A range of flowering times are found among forage and sweet sorghums [6]. Sweet sorghum genotypes with longer vegetative growth duration have larger stems that have greater potential for sucrose accumulation [6–8].

Flowering time is regulated by development, day length, phytohormones, shading, temperature, and the circadian clock [9–11]. In the long-day plant *Arabidopsis thaliana*, circadian and light signals are integrated to increase the expression of *FLOWERING LOCUS T (FT*) and flowering in long days. *FT* encodes a signaling protein synthesized in leaves that moves through the phloem to the shoot apical meristem (SAM) where it interacts with *FLOWERING LOCUS D (FD*) and reprograms the vegetative shoot apical meristem for reproductive development [12,13]. Expression of circadian clock genes such as *LATE ELONGATED HYPOCOTYL (LHY*) and *TIMING OF CAB1 (TOC1*) regulate the expression of the clock output gene *GIGANTEA (GI*) and genes in the flowering time pathway [14–16]. Photoperiod and circadian clock signals are integrated to control the expression and stability of CONSTANS (CO) an activator of *FT* expression [17]. Under inductive long day (LD) photoperiods, CO promotes the expression of *FT* which induces flowering in *Arabidopsis* [18].

Many of the genes in the *Arabidopsis* flowering time pathway are found in sorghum and other grass species such as *Oryza sativa* (rice) [10] and maize [19], however, the regulation of flowering time in these grasses has diverged from Arabidopsis in several important ways. Both rice and sorghum are facultative short-day (SD) plants. In rice, the expression of the **FT*-like* gene *Heading date 3a (Hd3a*) is promoted in SD [20]. In sorghum, expression of two different *FT*-like genes, *SbCN8* and *SbCN12*, is induced when plants are shifted from LD to SD [21,22]. In contrast to *Arabidopsis*, the rice and sorghum homologs of *CO* (rice *Heading date1, OsHd1; SbCO*) repress flowering in LD [10,23]. Rice and sorghum encode two additional grass-specific regulators of flowering Ehd1 and Ghd7. *Early heading date1* (Ehd1) activates the expression of *FT*-like genes, and *Grain number, plant height and heading date7* (Ghd7) represses the expression of *EHD1* and flowering [24,25]. When sorghum is grown in short days, SbEhd1 and SbCO induce the expression of *SbCN8* and *SbCN12*, leading to floral induction [21,22,26,27].

Under field conditions, time to flowering in sorghum varies from ~50 to >150 days after planting (DAP) depending on genotype, planting location and date (latitude/day-length), and the environment. A tall and “ultra-late” flowering sorghum variety called Milo Maize was introduced to the United States in the late 1800s [28]. Shorter and earlier flowering Milo genotypes such as Early White Milo and Dwarf Yellow Milo were selected from the introduced Milo genotype to promote improved grain yield in temperate regions of the US [1,28,29]. Genetic analysis determined that mutations in three independently segregating *Maturity (Ma*) loci (*Ma_1_, Ma_2_, Ma_3_*) were responsible for early flowering times in the Milo genotypes. A cross between Early White Milo (*ma1Ma_2_Ma_3_*) and Dwarf Yellow Milo (*Ma_1_ma_2_ma_3_*) was used to construct a set of Milo maturity standards, a series of nearly isogenic lines that differ at one or more of the *Maturity* loci (Quinby and Karper 1945, Quinby 1966, Quinby, 1967). A fourth *Maturity* locus (*Ma_4_*) was discovered in crosses of Milo (*Ma_4_*) and Hegari (*ma_4_*) [30]. More recent studies identified *Ma_5_* and *Ma_6_* [31]. Subsequent research showed that all of the Milos are dominant for *Ma_5_* and recessive for *ma_6_* [23,26]. In addition to these six *Ma* loci, many other flowering time quantitative trait loci (QTL) have been identified in sorghum [2,32–35]. Additional research has linked several of these QTL to genes such as *SbEHD1* and *SbCO* that are activators of *SbCN8* and *SbCN12* expression, sources of florigen in sorghum.

The genes corresponding to four of the six *Maturity* loci have been identified. *Ma_1_*, the locus with the greatest influence on flowering time photoperiod sensitivity, encodes *SbPRR37*, a pseudo-response regulator that inhibits flowering in LD [21]. *Ma_3_* encodes phytochome B [36], *Ma_5_* encodes phytochrome C [23], and *Ma_6_* encodes *Ghd7* a repressor of flowering in long days [26]. The genes corresponding to *Ma_2_* and *Ma_4_* have not been identified but recessive alleles at either locus results in early flowering in long days in genotypes that are photoperiod sensitive (*Ma_1_*) [28]. Prior studies also noted that genotypes recessive for *Ma_2_* flower later in genotypes that are photoperiod insensitive and recessive for *Ma_1_* and *Ma_6_* [28].

In this study, the impact of *Ma_2_* alleles on the expression of genes in the sorghum flowering time pathway was characterized. A QTL corresponding to *Ma_2_* was mapped and a candidate gene for *Ma_2_* identified by fine mapping and genome sequencing. The results show that *Ma_2_* enhances *Ma_1_ (SbPRR37*) and *SbCO* expression consistent with the impact of *Ma_2_* alleles on flowering time in genotypes that vary in *Ma_1_* alleles.

## Methods

### Plant growing conditions and populations

The cross of 100M and 80M was carried out by the Sorghum Breeding Lab at Texas A&M University in College Station, TX. F_1_ plants were grown in the field in Puerto Rico and self-pollinated to generate the F_2_ population used in this study. The 100M/80M F_2_ population was planted in the spring of 2008 at the Texas A&M Agrilife Research Farm in Burleson County, Texas (near College Station, TX).

The cross of Hegari and 80M was made in the greenhouse at Texas A&M University in College Station, TX. F_1_ plants were confirmed and self-pollinated to generate the F_2_ population used in this study. The Hegari/80M F_2_ population (n = 432) was planted in the spring of 2011 in the greenhouse in 18 L nursery pots in a 2:1 mixture of Coarse Vermiculite (SunGro Horticulture, Bellevue, WA) to brown pasture soil (American Stone and Turf, College Station, TX). All subsequent generations of Hegari/80M for fine mapping were grown in similar conditions. Greenhouse-grown plants were watered as needed and fertilized every two weeks using Peters general purpose 20-20-20 (Scotts Professional).

For circadian gene expression experiments, 100M and 80M genotypes were planted in MetroMix 900 (Sungro Agriculture) in 6 L pots, and thinned to 3 plants/pot after 2 weeks. Plants were grown in the greenhouse under 14 h days until 30 days after planting (DAP). After 30 days, the plants were moved into growth chambers and allowed to acclimate for 3 days. The growth chamber was set to 30°C and 14/10h L/D for the 3 days of entrainment and the first 24 h of tissue collection. The lights were changed to constant light for the second 24 h of tissue collection.

### QTL mapping and multiple-QTL analysis

DNA was extracted from leaf tissue for all individuals described above as described in the FastDNA Spin Kit manual (MP Biomedicals). All individuals in each mapping or HIF population were genotyped by Digital Genotyping using FseI digestion enzyme as described in Morishige et al [37]. DNA fragments were sequenced using the Illumina GAII platform and the reads were mapped back to the sorghum reference genome (v1.0, Phytozome v6). Genetic maps were created using MapMaker 3.0B with the Kosambi function [38]. QTL were mapped using WinQTLCartographer (v2.5.010) using composite interval mapping with a 1.0 cM walk speed and forward and backward model selection [39]. The threshold was set using 1000 permutations and α= 0.05. Upon release of v3.1 of the sorghum reference genome, the QTL coordinates were updated [40].

To look for possible gene interactions multiple-QTL analysis was used in the Hegari/80M F_2_ population. A single QTL analysis using the EM algorithm initially identified two primary additive QTL which were used to seed model selection. The method of Manichaikul et al. [41] was employed for model selection as implemented in R/qtl for multiple-QTL analysis [42]. Computational resources on the WSGI cluster at Texas A&M were used to calculate the penalties for main effects, heavy interactions, and light interactions. These penalties were calculated from 24,000 permutations for flowering time to find a significance level of 5% in the context of a two-dimensional, two-genome scan.

### Fine mapping of the *Ma_2_* QTL

All fine mapping populations for the *Ma_2_* QTL were derived from F2 individuals from the Hegari/80M population. The genetic distance spanning the *Ma_2_* locus is 2 cM corresponding to a physical distance of ~1.8 Mbp, so 1000 progeny would be required to obtain 20 recombinants within the *Ma_2_* QTL region. Six individuals that were heterozygous across the *Ma_2_* QTL were self-pollinated to generate six heterogeneous inbred families (HIFs) totaling 1000 F_3_ individuals. These individuals were grown out in the greenhouse, and flowering time was recorded. They were genotyped by Digital Genotyping as described above [37]. Two F_3_ individuals that had useful breakpoints with a heterozygous genotype on one side of the breakpoint were grown and self-pollinated to generate an additional round of HIFs (F_4_, n = 150) that were planted in the spring of 2013 and analyzed as described above. No new breakpoints were identified in the F_4_ generation, so this process was repeated again to generate F_5_ plants in the spring of 2014.

### Circadian gene expression analysis

For the circadian gene expression analysis, 30-day-old plants were placed in a growth chamber set to 14 h days for the first 24 h and constant light for the second 24 h at 30°C. Plants were entrained for 3 d before beginning tissue collection. Leaf tissue was collected and pooled from 3 plants every 3 h for 48 h. The experiment was repeated three times for a total of three biological replicates. RNA was extracted from each sample using the Direct-Zol™ RNA Miniprep Kit (Zymo Research) according to the kit instructions. cDNA was synthesized using SuperScript III kit for qRT-PCR (Invitrogen) according to the kit instructions. Primers for sorghum flowering pathway genes were developed previously, and primer sequences are available in Murphy et al [21]. Primer sequences for *Ma_2_* are available in S1 Table. Relative expression was determined using the comparative cycle threshold (Ct) method. Raw Ct values for each sample were normalized to C_t_ values for the reference gene *SbUBC* (Sobic.001G526600). Reference gene stability was determined previously [43]. ΔΔC_t_ values were calculated relative to the sample with the highest expression (lowest C_t_ value). Relative expression values were calculated with the 2^-ΔΔCt^ method [44]. Primer specificity was tested by dissociation curve analysis and gel electrophoresis of qRT-PCR products.

### Ma_2_ phylogenetic analysis

Protein sequences of the closest homologs of Ma_2_ were identified using BLAST analysis. Protein sequences were aligned using MUSCLE [45] and visualized using Jalview [46]. Evolutionary trees were inferred using the Neighbor-Joining method [47] in MEGA7 [48]. All positions containing gaps and missing data were eliminated.

### *Ma_2_* DNA sequencing and whole genome sequence analysis

Whole genome sequence reads of 52 sorghum genotypes including 100M and 80M were obtained from Phytozome v12. Base quality score recalibration, INDEL realignment, duplicate removal, joint variant calling, and variant quality score recalibration were performed using GATK v3.3 with the RIG workflow [49]. Sobic.002G302700 was sequenced via Sanger sequencing in the genotypes in Table 1 according to the BigDye Terminator Kit (Applied Biosystems). Primers for template amplification and sequencing are provided in S1 Table.

**Table 1.** Sequence variants of Sobic.002G203700 and their predicted effect on protein function

| Genotype | Historical *Ma<sub>2</sub>* allele | Sequence variant | Effect on protein function |
| --- | --- | --- | --- |
| 100M | *Ma<sub>2</sub>* | - | - |
| SM100 | *Ma<sub>2</sub>* | - | - |
| SM90 | *Ma<sub>2</sub>* | - | - |
| Hegari | *Ma<sub>2</sub>* | - | - |
| 80M | *ma<sub>2</sub>* | L141* | Deleterious |
| SM80 | *ma<sub>2</sub>* | L141* | Deleterious |
| 60M | *ma<sub>2</sub>* | L141* | Deleterious |
| 44M | *ma<sub>2</sub>* | L141* | Deleterious |
| 38M | *ma<sub>2</sub>* | L141* | Deleterious |
| 58M | *ma<sub>2</sub>* | L141* | Deleterious |
| Kalo | *ma<sub>2</sub>* | L141* | Deleterious |
| IS3614-2 | - | M83T | Deleterious |

\*Sequenced by Sanger sequencing

## Results

### Effects of *Ma_2_* alleles on flowering pathway gene expression

The recessive *ma_2_*-allele in 80M (*Ma_1_**ma_2_**Ma_3_Ma_4_Ma_5_ma_6_*) was previously reported to cause 80M to flower earlier than100M (*Ma_1_**Ma_2_**Ma_3_Ma_4_Ma_5_ma_6_*) in long days [28]. To help elucidate how *Ma_2_* modifies flowering time, we investigated the impact of *Ma_2_* alleles on the expression of genes in sorghum’s flowering time pathway. Gene expression was analyzed by qRT-PCR using RNA isolated from 100M (*Ma_2_*) and 80M (*ma_2_*) leaves collected every 3 hours for one 14h light/10h dark cycle and a second 24-hour period of constant light.

*SbPRR37* is a central regulator of photoperiod sensitive flowering in sorghum that acts by repressing the expression of *SbCN* (*FT*-like) genes in LD [21]. *SbPRR37* expression in 100M and 80M grown in long days peaked in the morning and again in the evening as previously observed [21] (Fig 1). The amplitude of both peaks of *SbPRR37* expression was reduced in 80M (*ma_2_*) compared to 100M (*Ma_2_*) (Fig 1A). *SbCO* also shows peaks of expression in the morning (dawn) and in the evening (~14h) [21] (Fig 5C). Analysis of *SbCO* expression in 100M and 80M showed that both peaks of *SbCO* expression were reduced in 80M compared to 100M (Fig 1B).

**Fig 1.**
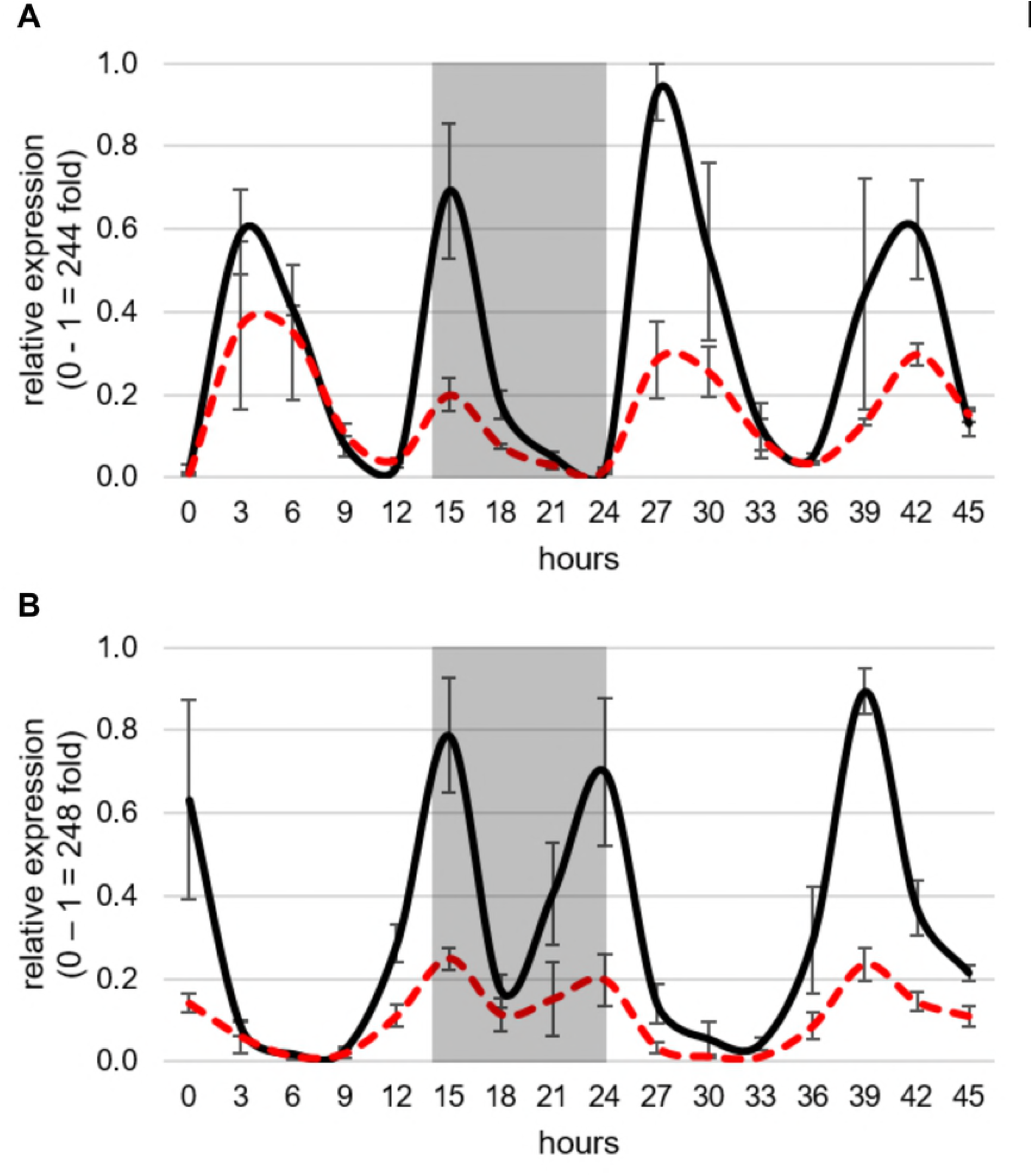
Circadian expression of genes regulating flowering in *S. bicolor* in 100M and 80M under long days. (A) Expression of *SbPRR37* in 100M (solid black lines) and 80M (dashed red lines). The expression peaks of *SbPRR37* are reduced in 80M. This is consistent with earlier flowering in 80M because *SbPRR37* represses the expression of the sorghum *FT*-like genes. (B) Expression of *SbCO* in 100M and 80M. Expression peaks of *SbCO* are also reduced in 80M. This is consistent with earlier flowering in 80M because under long days *SbCO* is a repressor of flowering. All expression values are normalized to *SbUBC* and are the mean of 3 biological replicates.

*SbCN8, SbCN12*, and *SbCN15* are homologs of *AtFT* that encode florigens in sorghum [22]. Expression of *SbCN8* and *SbCN12* increases when sorghum plants are shifted from LD to SD, whereas *SbCN15* shows minimal response to day length [21,26]. SbPRR37 and SbCO are co-repressor of the expression of *SbCN8* and *SbCN12* in long days, therefore, the influence of *Ma_2_* alleles on *SbCN8/12/15* expression was investigated [21,27]. When plants were grown in long days, expression of *SbCN12* was ~10 fold higher in 80M compared to 100M consistent with earlier flowering in 80M (Fig 2).

**Fig 2.**
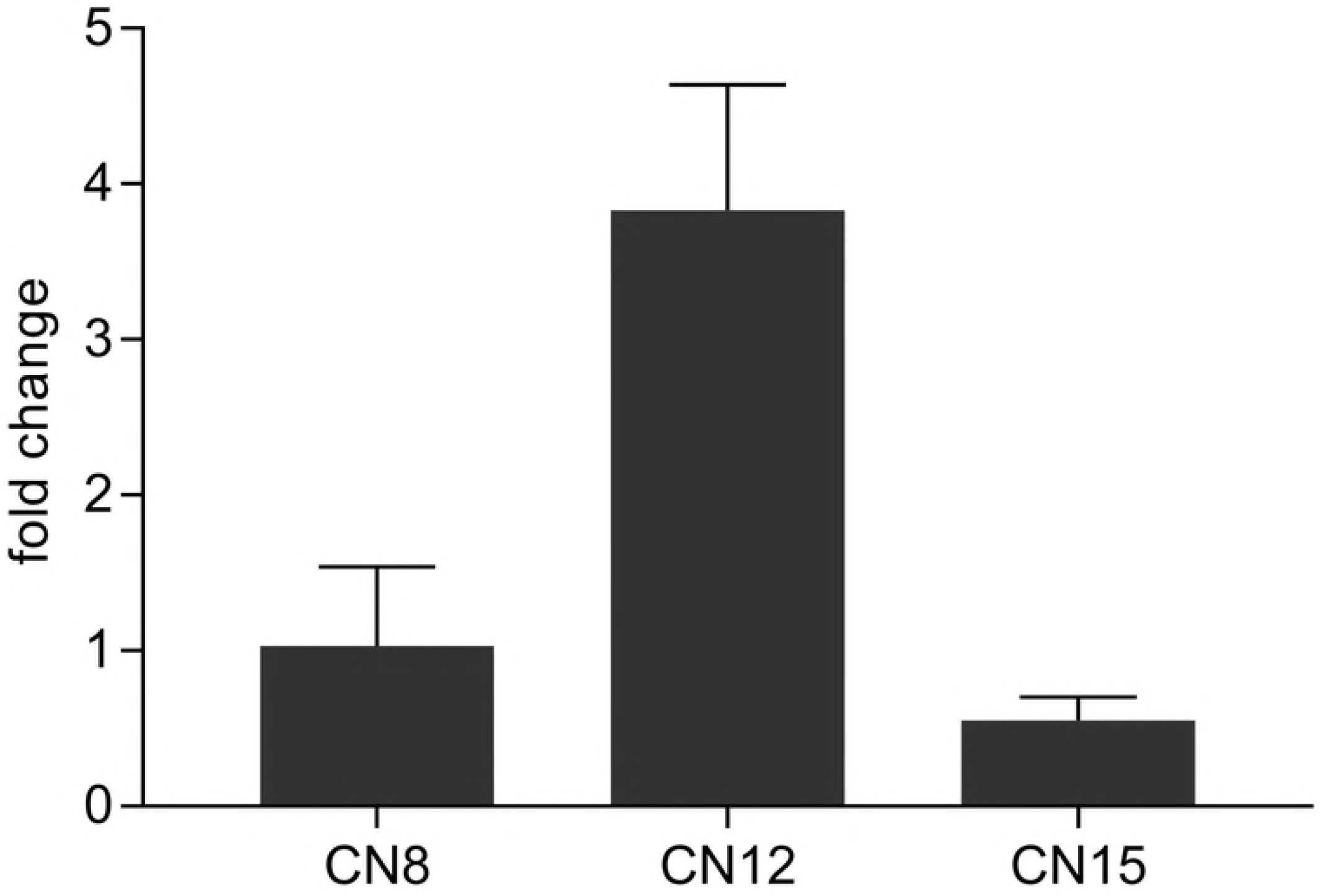
Expression of the *S. bicolor FT*-like genes *SbCN8, SbCN12*, and *SbCN15* in long days at the expected peak of expression. Expression of *SbCN* genes are all elevated in 80M, which is consistent with earlier flowering in that genotype. All expression values are normalized to *SbUBC* and are the mean of 3 biological replicates. Fold change was calculated as 2^-[Ct(100M)-Ct(80M)]^.

Previous studies showed that SbGHD7 represses *SbEHD1* expression and that alleles of *SbGHD7* differentially affect *SbCN8* expression (>SbCN12) [26]. Analysis of *SbEHD1* and *SbGHD7* expression in 100M and 80M showed that *Ma_2_* alleles modify the expression of these genes only to a small extent (S1 Fig).

The timing of the two daily peaks of *SbPRR37* and *SbCO* expression in sorghum is regulated by the circadian clock [21,26]. Therefore, it was possible that *Ma_2_* modifies *SbPRR37/SbCO* expression by altering clock gene expression. However, expression of the clock genes *TOC1* and *LHY* was similar in 100M and 80M (S1 Fig). Taken together, these results show that Ma_2_ is an activator of *SbPRR37* and *SbCO* expression in long days. Prior studies showed that co-expression of *SbPRR37* and *SbCO* in long days inhibits expression of *SbCN12* and floral initiation [27]. Later flowering in sorghum genotypes that are *Ma_1_Ma_2_* vs. *Ma_1_ma_2_* in long days is consistent with lower *SbCN12* expression in *Ma_1_Ma_2_* genotypes.

### Genetic analysis of Ma2 and Ma4

An F_2_ population derived from a cross of 100M (*Ma_2_*) and 80M (*ma_2_*) was generated to map the *Ma_2_* locus. Because 100M and 80M are nearly isogenic lines that differ at *Ma_2_*, only *Ma_2_* alleles were expected to affect flowering time in this population [28]. The F_2_ population (n = ~1100) segregated for flowering time in a 3:1 ratio as expected. The parental lines and F_2_ individuals were genotyped by Digital Genotyping (DG) which identifies single nucleotide polymorphism (SNP) markers in thousands of sequenced sites that distinguish the parents of a population [37]. The near isogenic nature of the parental lines resulted in a very sparse genetic map that lacked coverage of large regions of the sorghum genome including all of the long arm of SBI02. In retrospect, no *Ma_2_* QTL for flowering time was identified using this genetic map because the gene is located on the long arm of SBI02 (see below).

To overcome the limitations associated with the 80M/100M population, a second mapping population was created to identify the genetic locus associated with *Ma_2_*. An F_2_ population (n = 215) that would segregate for *Ma_2_* and *Ma_4_* was constructed by crossing Hegari (*Ma_1_**Ma_2_**Ma_3_**ma_4_**Ma_5_ma_6_*) and 80M (*Ma_1_**ma_2_**Ma_3_**Ma_4_**Ma_5_ma_6_*) [30,50]. The population was grown in a greenhouse under long day conditions and phenotyped for days to flowering. QTL for flowering time were identified on SBI02 and SBI10 (Fig 3). Recessive alleles of *Ma_2_* and *Ma_4_* result in earlier flowering when plants are grown in long days. The Hegari haplotype across the QTL on SBI10 was associated with early flowering therefore this QTL corresponds to *Ma_4_* (S3 Fig). The 80M haplotype across the QTL on SBI02 was associated with early flowering therefore the QTL on SBI02 corresponds to *Ma_2_*.

**Fig 3.**
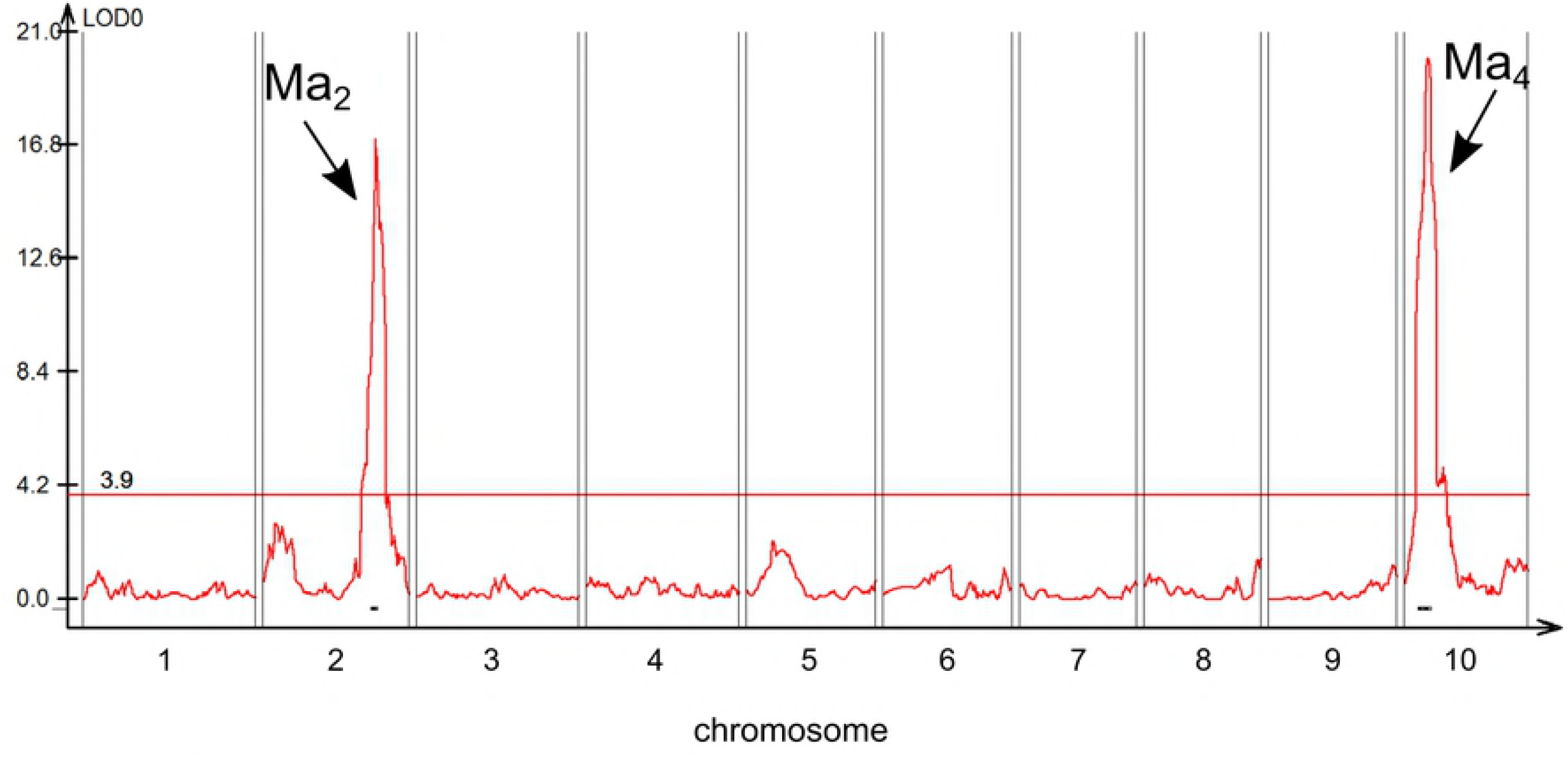
Quantitative trail locus (QTL) map of flowering time in the Hegari/80M F_2_ population. Two QTL were identified for variation in flowering time in the F2 population derived from Hegari (*Ma_1_Ma_2_Ma_3_ma_4_*) and 80M (*Ma_1_ma_2_Ma_3_Ma_4_*). This population was expected to segregate for *Ma_2_* and *Ma_4_*. Each recessive *Ma* allele causes earlier flowering. The QTL on LG10 corresponds to *Ma_4_* because F2 individuals carrying the Hegari allele contributed to accelerated flowering. F_2_ individuals carrying the 80M allele at the QTL on LG02 flowered earlier, so this QTL corresponds to *Ma_2_*.

### Epistatic interactions between *Ma_2_* and *Ma_4_*

Previous studies indicated an epistatic interaction exists between *Ma_2_* and *Ma_4_* [28]. Therefore, Multiple QTL Mapping (MQM) analysis [51] was employed, using data from the Hegari/80M F_2_ population, to identify additional flowering time QTL and interactions amongst the QTL as previously described [52]. MQM analysis identified the QTL for flowering time on SBI02 and SBI10 and an additional QTL on SBI09. Additionally, an epistatic interaction was identified between *Ma_2_* and *Ma_4_* (pLOD = 42). Interaction plots showed that in a dominant *Ma_4_* background, a dominant allele at *Ma_2_* delays flowering, while in a recessive *Ma_4_* background, *Ma_2_* has a minimal impact on flowering time (Fig 4). The interaction between *Ma_2_* and *Ma_4_* identified by MQM analysis is consistent previous observations that in a recessive *ma_4_* background flowering is early regardless of allelic variation in *Ma_2_* [28].

**Fig 4.**
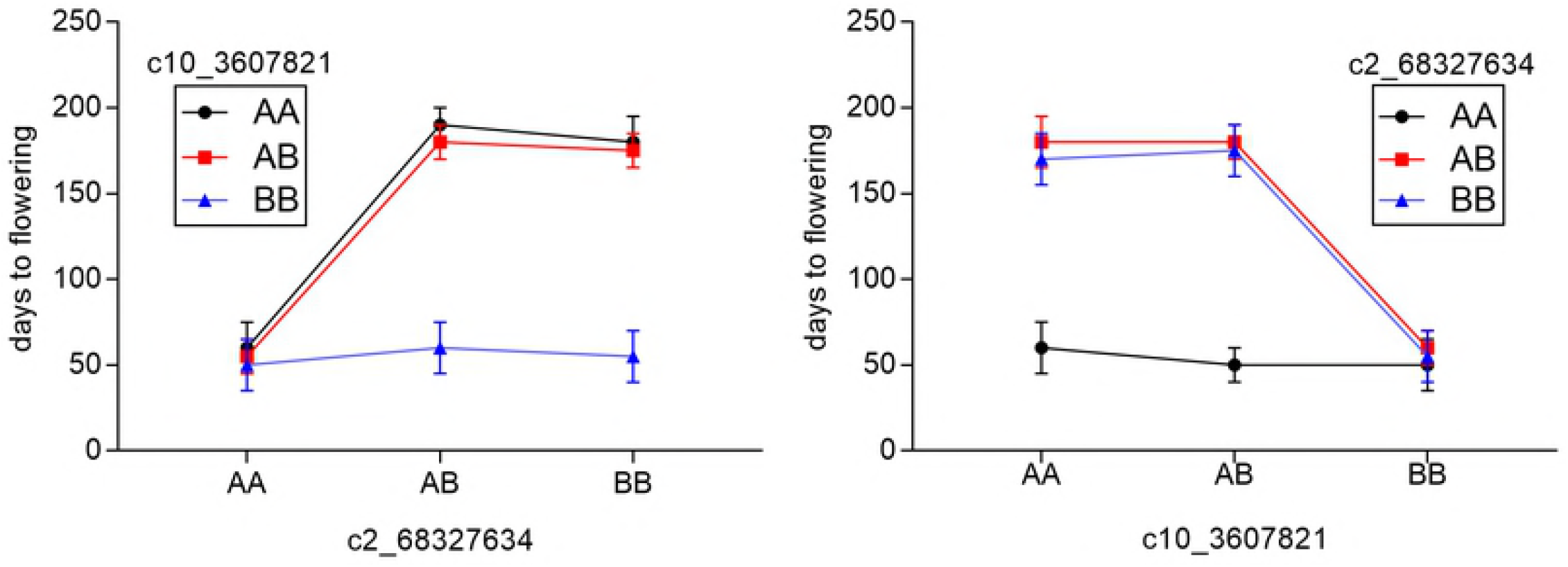
Interaction plots for the Ma2 QTL and the Ma4 QTL. There is a known interaction between *Ma_2_* (represented by marker c2_68327634) and *Ma_4_* (represented by marker c10_3607821). This interaction was identified by multiple QTL mapping (MQM). Dominant alleles of the *Ma* genes delay flowering. In a recessive ma_4_ background (AA at c2_68327634), the effect of *Ma_2_* on days to flowering is reduced. A represents the 80M allele and B represents the Hegari allele at each QTL. Reciprocal plots are shown.

### Ma2 candidate gene identification

The Hegari/80M F_2_ population located *Ma_2_* on SBI02 between 67.3 Mbp to 69.1 Mbp (Fig 5). To further delimit the *Ma_2_* locus, six lines from the Hegari/80M population that were heterozygous across the *Ma_2_* QTL but fixed across the *Ma_4_* locus (*Ma_4_Ma_4_*) were selfed to create heterogeneous inbred families (HIFs) (n=1000 F_3_ plants) [53]. Analysis of these HIFs narrowed the region encoding *Ma_2_* to ~600 kb (67.72 Mb-68.33 Mb) (Fig 5). Genotypes that were still heterozygous across the delimited locus were selfed and 100 F_4_ plants were evaluated for differences in flowering time. This process narrowed the *Ma_2_* locus to a region spanning ~500 kb containing 76 genes (67.72Mb-68.22Mb) (Fig 5, S2 Table).

**Fig 5.**
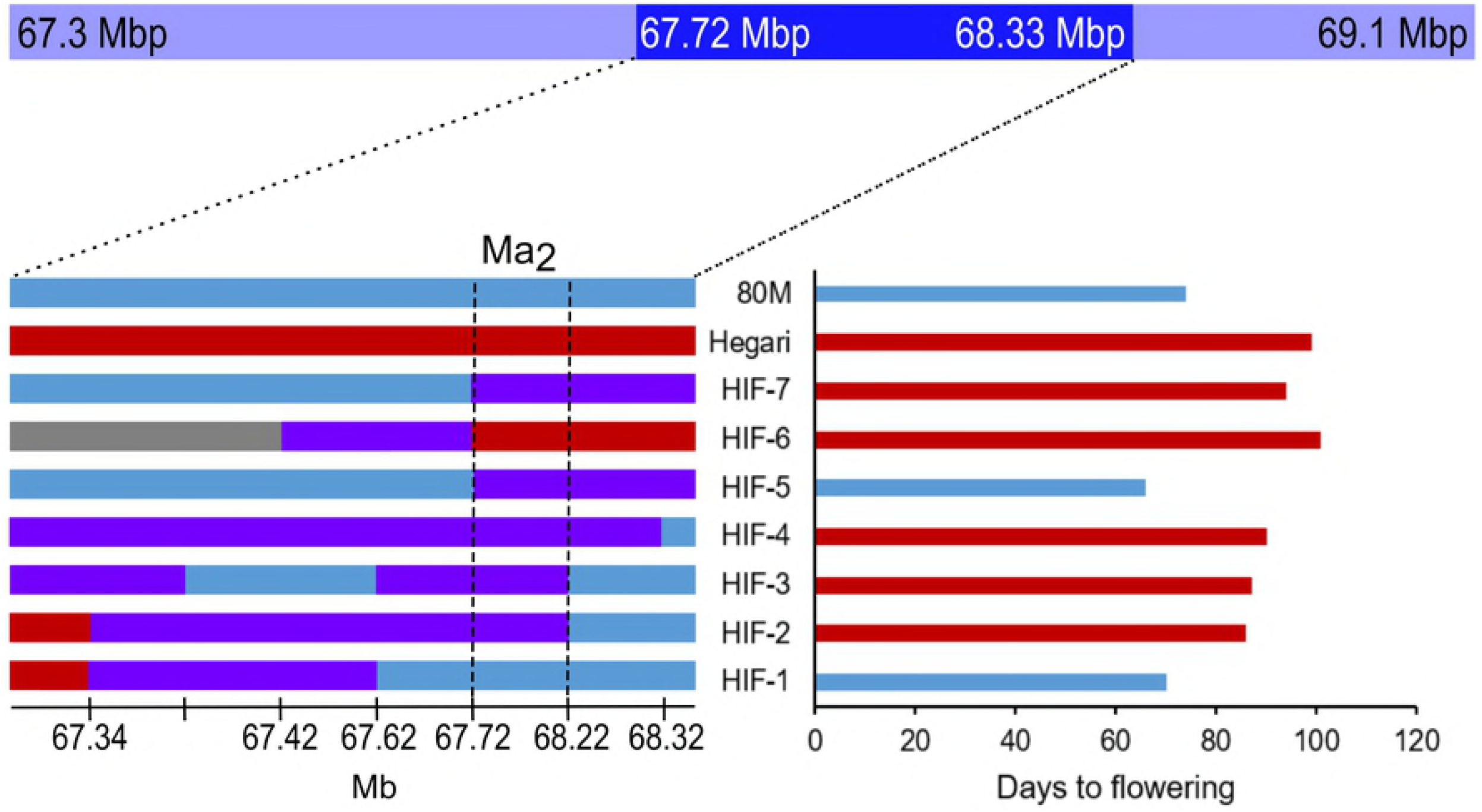
Fine-mapping of the Ma_2_ QTL. The *Ma_2_* QTL spans from 67.3 Mpb to 69.1 Mbp (light blue bar). Five F_2_ individuals that were heterozygous across the *Ma_2_* QTL were self-pollinated to generate heterogeneous inbred families (HIFs) totaling 1000 F_3_ individuals. Genotype and phenotype analysis of these HIFs narrowed the QTL region to ~600 kb (darker blue bar). Two additional rounds of fine-mapping narrowed the QTL region to ~500 kb (vertical dashed lines). This region contained 76 genes. The genotypes of relevant HIFs and the parents are shown to the left and their corresponding days to flowering are shown to the right. Blue regions correspond to the 80M genotype and red regions correspond to the Hegari genotype. Purple regions are heterozygous.

The low rate of recombination across the *Ma_2_* locus led us to utilize whole genome sequencing in conjunction with fine mapping to identify a candidate gene for *Ma_2_*. Since 100M and 80M are near isogenic lines that have very few sequence differences along the long arm of SBI02 where the *Ma_2_* QTL is located, whole genome sequences (WGS) of 100M and 80M were generated in collaboration with JGI (sequences available at www.phytozome.jgi.doe.gov). The genome sequences were scanned for polymorphisms within the 500 kb locus spanning *Ma_2_*. Only one T → A single nucleotide polymorphism (SNP) located in Sobic.002G302700 was identified that distinguished 100M and 80M within the region spanning the *Ma_2_* locus. The T → A mutation causes a Lys141* change in the third exon, resulting a truncated protein. A 500 bp DNA sequence spanning the T to A polymorphism in Sobic.002G302700 was sequenced from 80M and 100M to confirm the SNP identified by comparison of the whole genome sequences (Table 1). The T → A point mutation was present in 80M (*ma_2_*) whereas 100M (*Ma_2_*) encoded a functional version of Sobic.002G302700 that encodes a full length protein. Since this mutation was the only sequence variant between 100M and 80M in the fine-mapped locus, Sobic.002G302700 was identified as the best candidate gene for *Ma_2_*.

Sobic.002G302700 is annotated as a SET (Suppressor of variegation, Enhancer of Zeste, Trithorax) and MYND (Myeloid-Nervy-DEAF1) (SMYD) domain-containing protein. SMYD domain family proteins in humans have been found to methylate histone lysines and non-histone targets and have roles in regulating chromatin state, transcription, signal transduction, and cell cycling [54,55]. The SET domain in SMYD-containing proteins is composed of two sub-domains that are divided by the MYND zinc-finger domain. The SET domain includes conserved sequences involved in methyltransferase activity including nine cysteine residues that are present in the protein encoded by Sobic.002G303700 (Fig 6) [56]. The MYND domain is involved in binding DNA and is enriched in cysteine and histidine residues [57]. Protein sequence alignment of Sobic.002G302700 homologs revealed that the SYMD protein candidate for Ma_2_ is highly conserved across flowering plants (Fig 6).

**Fig 6.**
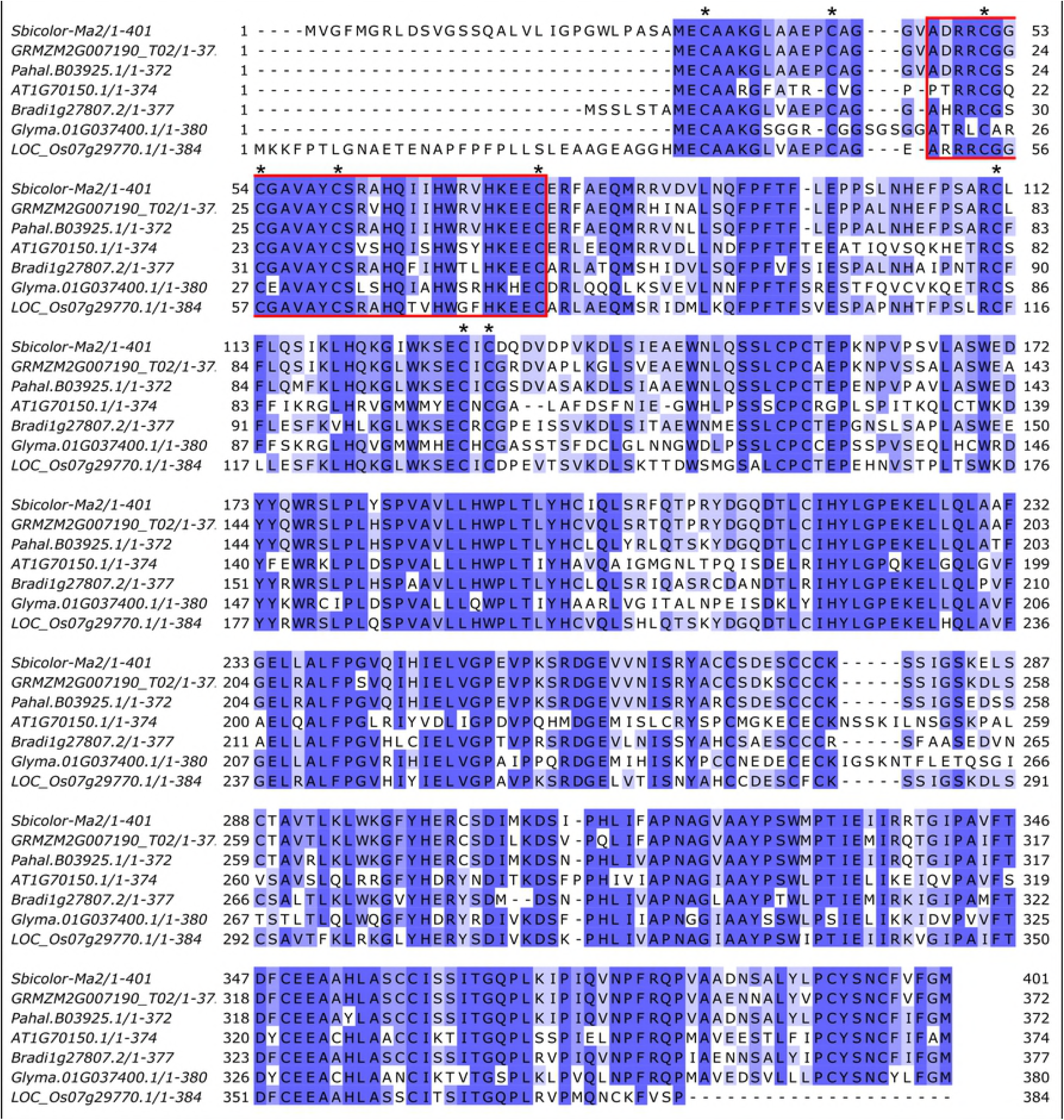
Alignment of Sobic.002G302700 with its closest homologs in several plant species. Sobic.002G302700 is highly conserved across plant species. It is annotated as a Set and MYND (SMYD) protein. SMYD proteins have lysine methyltransferase activity. The MYND region is highlighted in red. The nine conserved Cys residues typical of SMYD proteins are indicated by asterisks.

To learn more about *Ma_2_* regulation, the expression of Sobic.002G302700 in 100M and 80M was characterized during a 48h L:D/L:L cycle. *Ma_2_* showed a small increase in expression from morning to evening and somewhat higher expression in 100M compared to 80M during the evening (S1 Fig).

### Distribution of Ma_2_ alleles in the sorghum germplasm

Recessive *ma_2_* was originally found in the Milo background and used to construct Double Dwarf Yellow Milo (*Ma_1_**ma_2_**ma_3_Ma_4_Ma_5_ma_6_*) [28]. Double Dwarf Yellow Milo was crossed to Early White Milo (*ma_1_**Ma_2_**Ma_3_Ma_4_Ma_5_ma_6_*) and the progeny selected to create 100M, 80M and the other Milo maturity standards [1,28,58]. Several of the Milo maturity standards were recorded as recessive *Ma_2_* (80M, 60M, SM80, SM60, 44M, 38M) and others as *Ma_2_* dominant (100M, 90M, SM100, SM90, 52/58M). In order to confirm the *Ma_2_* genotype of the maturity standards, the 500 bp sequence spanning the Lys141* mutation in Sobic.002G302700 was obtained from most of these genotypes (Table 1). Kalo was also identified as carrying a recessive allele of *Ma_2_*. Kalo was derived from a cross of Dwarf Yellow Milo (*ma_2_*), Pink Kafir (*Ma_2_*),and CI432 (*Ma_2_*), therefore it was concluded that DYM is the likely source of recessive *ma_2_* [28]. Sequence analysis showed that the genotypes previously identified as *ma_2_* including Kalo, 80M, SM80, 60M, 44M, 38M, and 58M carry the recessive mutation in Sobic.002G302700 identified in 80M. 100M, SM100, and Hegari that were identified as *Ma_2_*, did not contain the mutated version of Sobic.002G302700 (Table 1). Additionally, sequences of *Ma_2_* from 52 sorghum genotypes with publicly available genome sequences were compared [40]. Sobic.002G302700 was predicted to encode functional proteins in all except one of these sorghum genotypes. A possible second recessive *Ma_2_* allele was found in IS3614-2 corresponding to an M83T missense mutation that was predicted to be deleterious by PROVEAN [59].

## Discussion

Sorghum is a facultative short day plant. In photoperiod sensitive sorghum genotypes, following the vegetative juvenile phase, day length has the greatest impact on flowering time under normal growing conditions. The development of early flowering grain sorghum adapted to temperate regions of the US was based on the selection of mutations in numerous genes that reduced photoperiod sensitivity. Genetic analysis of the loci and genes containing these mutations beginning in the 1940’s [50,58] identified six *Maturity* loci (*Ma_1_-Ma_6_*) that resulted in earlier flowering time when plants were grown in long days. Recessive alleles at each of the six *Ma* loci reduces photoperiod sensitivity [30,31,58]. Molecular identification of the genes corresponding to *Ma_1_, Ma_3_, Ma_5_* and *Ma_6_* and other genes in the sorghum flowering time pathway (i.e., *SbCO, SbEHD1, SbCN8/12*) and an understanding of their regulation by photoperiod and the circadian clock led to the model of the flowering time pathway shown in Figure 7 [60]. The current study showed that Ma_2_ represses flowering in long days by increasing the expression of SbPRR37 (*Ma_1_*) and *SbCO*. The study also located QTL for *Ma_2_* and *Ma_4_*, confirmed an epistatic interaction between *Ma_2_* and *Ma_4_*, and identified a candidate gene for *Ma_2_*.

**Fig 7.**
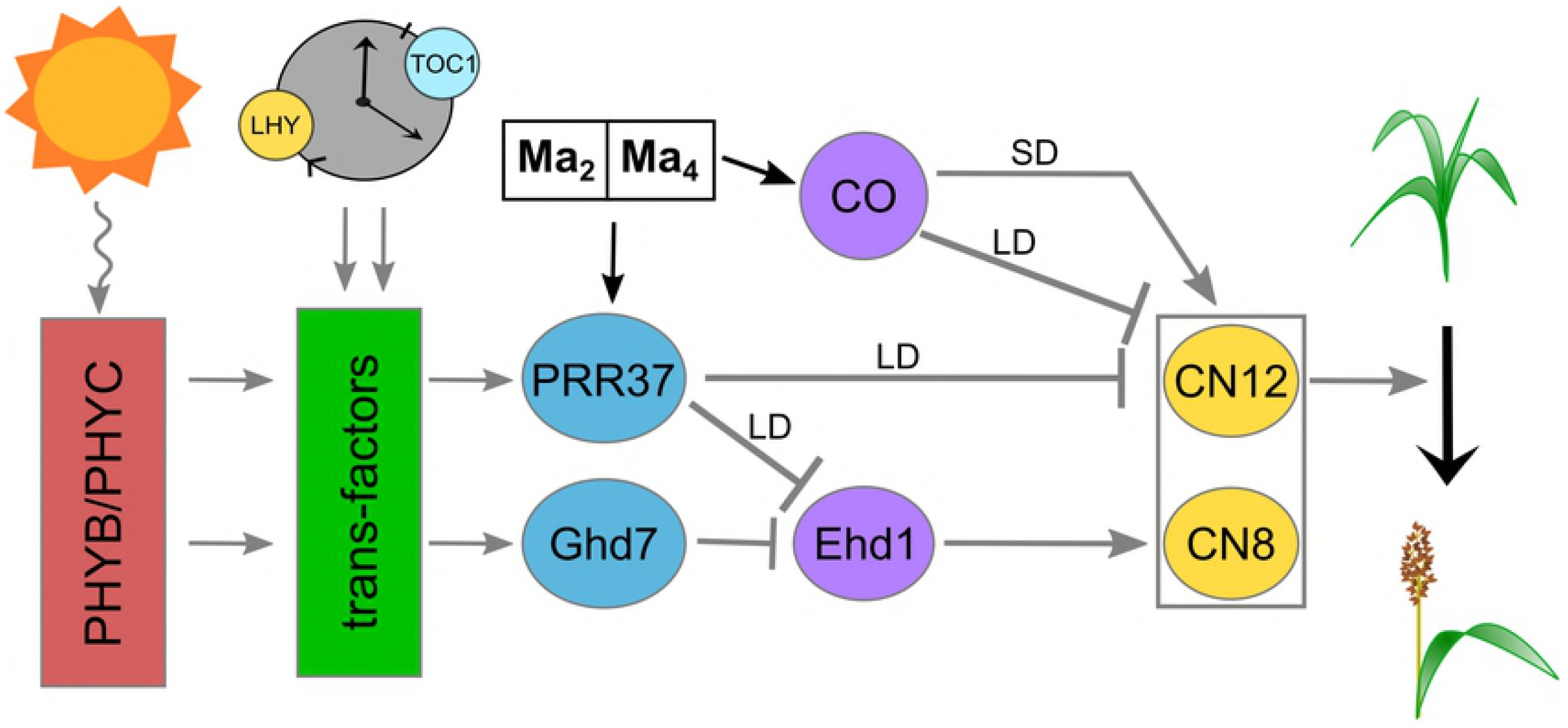
A model of the flowering time regulatory pathway in *S. bicolor*. *Ma_2_* and *Ma_4_* work codependently to enhance the expression of *SbPRR37* and *SbCO*. In LD, SbPRR37 and SbCO in turn repress the expression of the *SbCN* genes, especially *SbCN12*, to repress the floral transition.

The recessive *ma_2_* allele characterized in this study arose in a highly photoperiod sensitive Milo genotype that was introduced into the US in the late 1800’s and then selected for early flowering to enhance grain production. Quinby and Karper [29] created near isogenic Milo maturity genotypes with allelic variation at specific *Ma* loci to facilitate genetic and physiological analysis of flowering time regulation. In the current study, we utilized two of these maturity genotypes, 100M (*Ma_1_**Ma_2_**Ma_3_Ma_4_Ma_5_ma_6_*) and 80M (*Ma_1_**ma_2_**Ma_3_Ma_4_Ma_5_ma_6_*), to characterize how allelic variation in *Ma_2_* affects the expression of genes in the sorghum photoperiod regulated flowering time pathway (Fig 7). This analysis showed that mutation of *ma_2_* (80M) significantly reduced the amplitude of the morning and evening peaks of *SbPRR37* and *SbCO* expression compared to 100M (*Ma_2_*) without altering the timing of their expression. In addition, the expression of *SbCN12* (*FT*-like) increased 8-fold in leaves of 80M compared to 100M consistent with earlier flowering in 80M. In contrast, expression of clock genes (*TOC1, LHY*) and other genes (i.e., GHD7, EHD1) in the photoperiod regulated flowering time pathway were modified to only a small extent by allelic variation in *Ma_2_*. Based on these results, we tentatively place *Ma_2_* in the flowering time pathway downstream of day length sensing phytochromes and circadian clock regulation and identify *Ma_2_* as a factor that enhances *SbPRR37* and *SbCO* expression (Fig 7).

The differential increase in *SbCN12* expression in 80M (vs. 100M) is consistent with differential inhibition of *SbCN12* expression in long days by the concerted action of SbPRR37 and SbCO which has been previously shown to inhibit *SbCN12* expression [27]. Prior studies showed that 100M (*Ma_2_*) flowers later than 80M (*ma_2_*) in long days [28]. The impact of *Ma_2_* alleles on the expression of *SbPRR37* and *SbCO* is consistent with the effect of these alleles on flowering time in long days. Genetic studies showed that floral repression mediated by SbPRR37 requires SbCO as a co-repressor [27]. Therefore, enhanced expression of both *SbPRR37* (*Ma_1_*) and *SbCO* by *Ma_2_* in *Ma_1_Ma_2_* genotypes in long days is consistent with delayed flowering under these conditions relative to genotypes such as 80M that are *Ma_1_ma_2_*. Molecular genetic studies also showed that SbCO is an activator of *SbCN12* expression and flowering in *ma_1_* genetic backgrounds [27]. This is consistent with the observation that *ma_1_Ma_2_* genotypes flower earlier than *ma1ma_2_* genotypes when grown in long days [28].

### Interactions between *Ma_2_* and *Ma_4_*

Multiple QTL (MQM) analysis of results from a population derived from Hegari/80M identified an interaction between *Ma_2_* and *Ma_4_* as well as one additional flowering QTL on SBI09. Flowering time QTL on SBI09 have been identified in other mapping populations, but the gene(s) involved have not been identified [33,34]. The interaction between *Ma_2_* and *Ma_4_* confirmed previous observations that recessive *ma_4_* causes accelerated flowering in long days in *Ma_1_Ma_2_* genotypes [28]. Interestingly, the influence of *Ma_2_* and *Ma_4_* alleles on flowering time is affected by temperature [28,61]. The influence of temperature on flowering time pathway gene expression in 80M and 100M in the current study was minimized by growing plants at constant 30C. However, analysis of the temperature dependence of Ma_2_ and Ma_4_ on flowering time may help elucidate interactions between photoperiod and flowering time that have been previously documented [28,62]. Positional cloning of *Ma_4_* is underway to better understand the molecular basis of *Ma_2_* and *Ma_4_* interaction and their impact on flowering time.

### Identification of a candidate gene for *Ma_2_*

A mapping population derived from Hegari/80M that segregated for *Ma_2_* and *Ma_4_* enabled localization of the corresponding flowering time QTL in the sorghum genome (SBI02, *Ma*_2_; SBI10, *Ma_4_*). The *Ma_2_* QTL on SBI02 was fine-mapped using heterozygous inbred families (HIFs) from Hegari/80M. Several rounds of fine-mapping delimited the QTL to a ~500kb region containing 76 genes. Low recombination rates in this region of SBI02 made it difficult to delimit the QTL further using break point analysis therefore comparison of genome sequences from 80M and 100M was used to help identify a candidate gene for *Ma_2_*. The recessive *ma_2_* allele present in 80M arose in a Milo genotype similar to 100M [28] and genetic analysis of 100M and 80M showed that these near isogenic genotypes lacked DNA markers on the long arm of SBI02 where *Ma_2_* is located. Indeed, a scan of the whole genome sequences of 100M and 80M identified only a single T to A mutation in the 500 kb region spanning the fine-mapped *Ma_2_* locus. This mutation caused a Lys141* change in the third exon of Sobic.002G302700 resulting in protein truncation. Based on this information Sobic.002G302700 was tentatively identified as the best candidate gene for *Ma_2_*.

Sobic.002G302700 encodes a SET (Suppressor of variegation, Enhancer of Zeste, Trithorax) and MYND (Myeloid-Nervy-DEAF1) (SMYD) domain containing protein. In humans, SMYD proteins act as lysine methyltransferases, and the SET domain is critical to this activity. Therefore, Ma_2_ could be altering the expression of *SbPRR37* and *SbCO* by modifying histones associated with these genes. The identification of this SMYD family protein’s involvement in flowering in sorghum as well as the identification of highly conserved homologs in other plant species suggests that *Ma_2_* may correspond to a novel regulator of sorghum flowering. While a role for SYMD-proteins (lysine methyltransferases) as regulators of flowering time has not been previously reported, genes encoding histone lysine demethylases (i.e., JMJ30/32) have been found to regulate temperature modulated flowering time in Arabidopsis [63].

J.R. Quinby [50] identified only one recessive allele of *Ma_2_* among the sorghum genotypes used in the Texas sorghum breeding program. The maturity standard lines including 80M that are recessive for *ma_2_* and the genotype Kalo were reported to be derived from the same recessive *ma_2_* Milo genotype [28]. To confirm this, *Ma_2_* alleles in the relevant maturity standards and Kalo were sequenced confirming that all of these *ma_2_* genotypes carried the same mutation identified in 80M (Table 1). Among the 52 sorghum genotypes with available whole genome sequences, only 80M carried the mutation in Ma_2_ [40]. One possible additional allele of *ma_2_* was identified in IS36214-2, which contained a M83T missense mutation that was predicted to be deleterious to protein function by PROVEAN [59].

In conclusion, we have shown that *Ma_2_* represses flowering in long days by promoting the expression of the floral repressor *SbPRR37* and *SbCO*, a gene that acts as a co-repressor in long days (Fig 7). Sobic.002G302700 was identified as the best candidate for the sorghum *Maturity* locus *Ma_2_* although further validation such as targeted mutation of Sobic.002G302700 in a *Ma_1_Ma_2_* sorghum genotype or complementation of *Ma_1_ma_2_* genotypes will be required to confirm this gene assignment. The identification of this gene and its interaction with *Ma_4_* begins to elucidate a new element of the photoperiod flowering regulation pathway in sorghum.

## Acknowledgements

This research was supported by the Perry Adkisson Chair in Agricultural Biology and by the Agriculture and Food Research Initiative Competitive Grant 2016-67013-24617 from the USDA National Institute of Food and Agriculture. The authors would like to thank Dr. Daryl Morishige for his assistance in performing the Hegari/80M cross as well as Robin Poncik for her assistance in recording flowering dates.

## Supporting information

**S1 Fig. Circadian expression of Sobic.002G302700 in 100M and 80M**

The expression of Sobic.002G302700 does not cycle diurnally in 100M (solid black line) or 80M (dashed red line). There was no difference in expression between 100M and 80M in the first day. Expression was slightly elevated in 100M compared to 80M during the night and through the following morning.

**S2 Fig. Circadian expression of *SbTOC1, SbLHY, SbGhd7*, and *SbEhd1***

There were no consistent differences in expression of (A) *SbTOC1*, (B) *SbLHY*, (C) *SbGhd7*, and (D) *SbEhd1* between 100M (solid black line) and 80M (dashed red line).

**S3 Fig. Genotype x phenotype plots for the QTL on SBI02 and SBI10**

Recessive alleles of *Maturity* genes contribute to earlier flowering. 80M (AA) is recessive for *ma_2_*, while Hegari (BB) is dominant. Individuals genotyped AA for the QTL on SBI02 (represented by marker c2_68327634) flowered ~100 d earlier than those genotyped BB. 80M is dominant for *Ma_4_*, and individuals genotyped AA at the QTL on SBI10 (represented by marker c10_3607821) flowered ~100 d earlier than those genotyped BB.

**S1 Table. *Ma_2_* (Sobic.002G302700) sequencing and qPCR primers**

**S2 Table. Genes in the fine-mapped *Ma_2_* QTL region**

